# Tracking the phenology of riverine insect communities using environmental DNA

**DOI:** 10.1101/2025.01.10.632357

**Authors:** Eva Cereghetti, Xhesida Ajvazi, François Keck, Andrea Patrignani, Nicolò Tartini, Florian Altermatt, Luca Carraro

## Abstract

Aquatic insects are iconic and ecologically highly relevant inhabitants of riverine ecosystems. They are also often the target of monitoring programs to assess the ecological status of these lotic habitats. Environmental DNA (eDNA) techniques have been widely and successfully implemented to investigate freshwater insects and other macroinvertebrates. Commonly, such monitoring is conducted at one or two timepoints per year, despite the known strong seasonality and phenology of aquatic insects. Here, we assessed if and how eDNA can capture the temporal changes of the orders Ephemeroptera (mayflies), Plecoptera (stoneflies), Trichoptera (caddisflies) and Diptera (true flies). We carried out eDNA sampling at roughly monthly intervals from April to October at 25 sites across a whole river catchment in the northeastern part of Switzerland. We found pronounced, cyclic phenological trends in all orders but Trichoptera: the communities diverged from spring to summer and then in fall gradually returned closer to the spring state. The four orders exhibited different predominance in gains or losses of species detection throughout this time interval. Lastly, we found that field replicates, despite showing a relatively high local stochasticity, were able to provide a more complete assessment of aquatic communities and could thus be used as a proxy for the frequency of observation of a species through the seasons. In fact, this approach yielded comparable temporal patterns to the ones extracted from the Global Biodiversity Information Facility (GBIF). Overall, our findings demonstrate that eDNA techniques can be used to reveal intra-annual dynamics of aquatic insects. Given the current necessity to assess and monitor the biodiversity status of ecosystems, we therefore show that eDNA methods are a viable option to obtain a deeper understanding of the structuring of freshwater communities over time.

## INTRODUCTION

The widely reported dramatic decline of biodiversity across rivers globally (Dudgeon et al. 2006, Vörösmarty et al. 2010, WWF 2024) has prompted an increasing interest in assessing and monitoring the biodiversity of freshwaters to evaluate their current and future ecological integrity (Tickner et al. 2020, Sumudumali and Jayawardana 2021). The sampling of environmental DNA (eDNA)—that is, the extraction of organismal DNA from environmental substrates—and subsequent species assignation via metabarcoding has proved to be a viable tool to assess the composition of stream communities (Deiner et al. 2017, Altermatt et al. 2020), and it has to an extent demonstrated its capability of capturing seasonal signals in community changes (Jensen et al. 2021, Blackman et al. 2022, Sander et al. 2024). However, many freshwater monitoring programs and studies utilizing eDNA methods tend to sample only once or a few times a year, or have to concede spatial replication for a higher temporal resolution (Sander et al. 2024). Investigating intra-annual dynamics at the full scale of catchments can be fundamental to address the strong phenological signature that distinguishes many aquatic communities and can shed light on how the timing of sampling can influence findings (Šporka et al. 2006, Perry et al. 2024).

Freshwater systems are habitats that host an astoundingly high number of species compared to the small surface that they occupy (Dudgeon et al. 2006, Biggs et al. 2017). Aquatic insects are a central part of this diversity (Dijkstra et al. 2014): they fulfil a range of roles within not only the aquatic food web (Wallace and Webster 1996), but also the terrestrial food web (Gücker et al. 2009), and the sensitivity of some groups to environmental conditions has resulted in taxa being used as indicator species to determine the ecological status of the target systems (Barbour et al. 1999, Buss et al. 2014). Many of these aquatic insects display a strong phenology and have a life history development split between aquatic and terrestrial stages. Consequently, communities of aquatic macroinvertebrates can undergo substantial shifts throughout the year, driven by changes in the abundance and growth of aquatic larvae and by differences in the timing of their emergence (e.g., Cowell et al. 2004, Haidekker and Hering 2008, Wilmot et al. 2021). Such changes can result in altered species interaction and co-existence patterns (Rudolf 2019), which can further cascade to influence food webs and ecosystem functions (McMeans et al. 2015, Kortsch et al. 2021). Importantly, strong phenological dynamics can hamper the comparisons of samples taken at different times through the year and could confound ecological assessments relying on the presence of these organisms (Šporka et al. 2006, Esmaeili Ofogh et al. 2023). Despite the rise in popularity of eDNA and metabarcoding methods, only a few studies have investigated the capability of these techniques to reveal intra-annual dynamics of whole river catchments (Perry et al. 2024).

Yet, the use of eDNA is particularly suited to the investigation of aquatic insects within riverine networks. The downstream water flow in these systems behaves like a ‘conveyer belt’ of organismal material (Deiner et al. 2016), allowing eDNA sampling to present an integrated spatial signal of river biodiversity (Altermatt et al. 2020, Carraro et al. 2020). Furthermore, the development of targeted primers (Leese et al. 2021) has enabled accurate species level identification of a range of aquatic insects, which would otherwise require time consuming and challenging morphological identification (Brantschen et al. 2021). Environmental DNA methods come with their own challenges and should not be considered a direct substitute for more traditional methods (Pereira et al. 2021, Keck et al. 2022). However, past work has shown that eDNA can deliver a robust overview of community changes over short time intervals (Bista et al. 2017, Jensen et al. 2022, Sander et al. 2024). It thus represents a promising venue for investigating temporal dynamics in riverine systems and for revealing phenological signatures in aquatic insect communities.

In this study, we investigated the temporal distribution of Ephemeroptera, Plecoptera, Trichoptera and Diptera within the catchment of a temperate river system. We carried out regular (every 4 weeks) collections of eDNA samples over the span of seven months, with an extensive sampling site selection that aimed at maximizing coverage of the whole river system, and a large number of replicate samples collected per each site and time point, in a bid to unravel the uncertainty associated with single eDNA measurements. Specifically, we analyzed different facets of the temporal dynamics of the four insect groups, with the important assumption that the observed community changes were primarily driven by changes in the detectability of species rather than by extinction or colonization events. Our expectations were as follows. First, the variability in community composition detected between replicated eDNA samples collected at the same sampling site and time point should be considerably lower than between samples collected at different sites and/or time points. Second, our findings should reflect the seasonal ecology of aquatic insects and thus reveal temporal changes in the community driven by insect population dynamics and emergence patterns. Third, the timing and magnitude of these changes in detection should differ across the four orders due to the abundance and richness of these aquatic insects often showing asynchronous peaks across the orders (e.g., Waringer 1996, Haidekker and Hering 2008). Lastly, temporal changes in how frequently species were detected (i.e., flagged as present) during our eDNA campaign should match the reported frequencies of observations in the region, as found in the Global Biodiversity Information Facility (GBIF). If these expectations were met, eDNA could be considered a suitable method to assess phenological signatures of aquatic insect communities.

## METHODS

Our study system was the Necker river, draining an area of 126 km^2^ in the northeastern part of Switzerland (Figure 1). The upper part of the Necker flows at about 1150 m a. s. l., whereas its lowest point is located at 550 m a. s. l., where it joins the Thur river. Approximately a third of the catchment is covered by forests, while the rest is mostly covered by extensive agricultural land and barren area. The river remains relatively pristine, with minimal urban alterations. Overall, the Necker has a high diversity of invertebrates, as revealed by previous eDNA campaigns (Mächler et al. 2019, Carraro et al. 2020, 2023, Blackman et al. 2022).

**Figure 1.**
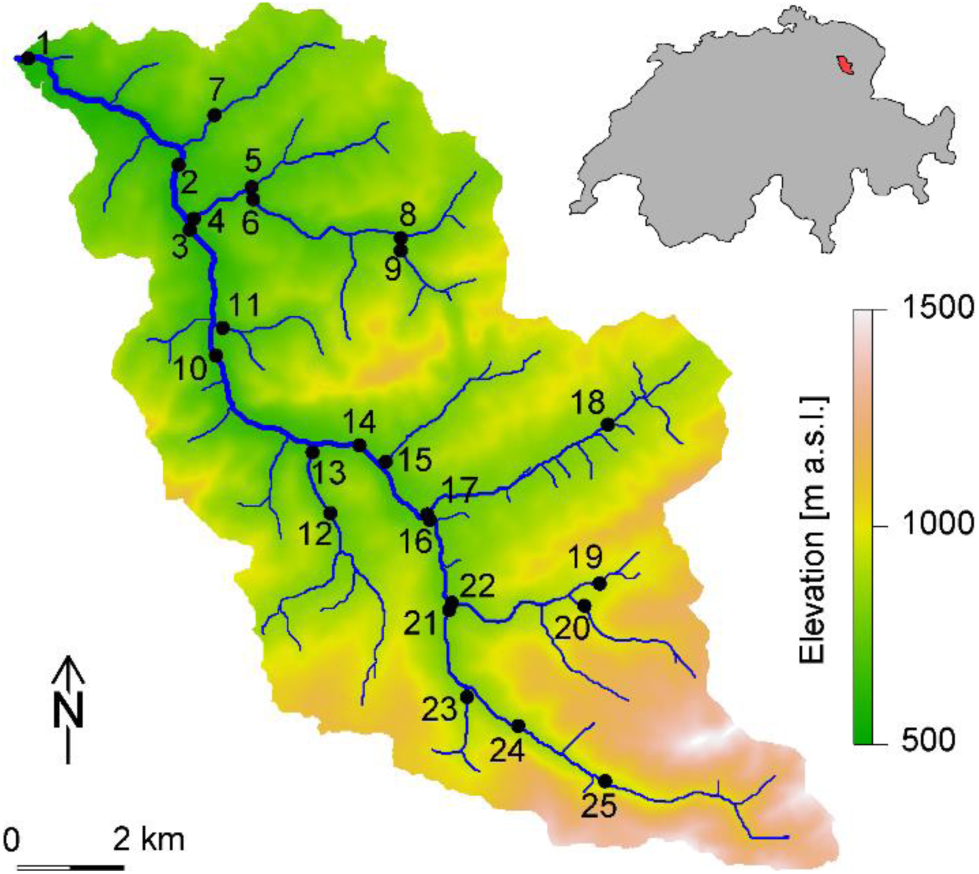
Map of the Necker catchment, located in the northeastern part of Switzerland (see inset on the top-right corner), and overview of the 25 sampled sites. The map was generated using the *rivnet* package (Carraro 2023).

We collected water samples from 25 sites along both main and side stems of the Necker to maximize coverage of the system (Figure 1). The sites were sampled eight times between April and October 2022 with a time interval of about four weeks between sampling bouts. Exact sampling dates were chosen by ensuring that water discharge was sufficiently close to its baseflow level, thus allowing an unbiased and practically feasible eDNA collection. Each sampling campaign took 2 to 3 days to collect samples from all 25 sites, and will hereafter be identified by its respective starting date: April 11, May 9, June 14, July 5, August 3, August 29, September 22, October 26.

For each site and time point, we used a peristaltic pump (Alexis Peristaltic Pump v2.0, Proactive Environmental Products, Florida, USA), to filter river water from the water column through a 0.22-μm Sterivex filter (Merck Millipore, Merck KgaA, Darmstadt, Germany). For each filter, the volume of water filtered ranged from 0.5 L (with more turbid waters leading to early filter clogging) to 1 L (in the case of clear waters). We repeated the process five times at each site to obtain five field replicates, thus totaling 1000 samples across the whole study campaign (25 sites × 8 time points × 5 field replicates). Twice per sampling bout, we filtered UV-treated Nanopure water as field negative controls (n = 16). Filters were stored in a cool box filled with ice after collection in the field, then transported to the laboratory and preserved at −20 °C until further processing.

### DNA extraction

We carried out the DNA extraction and PCR set up in a specialized clean lab facility at Eawag, Switzerland (Deiner et al. 2015). Prior to extraction, we defrosted all Sterivex filters at room temperature for 15 minutes and wiped them with a 30% bleach solution to remove any DNA possibly remaining on the filter external surface. We disinfected the laboratory bench with a 30% bleach solution followed by wiping with 70% ethanol, whereas we exposed pipettes and vortex table to UV radiation for a minimum of 20 minutes prior to use. We extracted environmental DNA using the DNeasy PowerWater Sterivex kit (Qiagen, Hilden, Germany) and following the kit’s provided protocol, except for skipping the mechanical lysis step as eDNA was not encapsulated in a membrane. Alongside the samples, we extracted 40 Sterivex filters through which we had run MiliQ water, and which served as extraction negative control. Samples were extracted in randomized order after each field sampling time point then stored at −20 °C.

### Library preparation and sequencing

DNA metabarcoding was performed by targeting a 142-base pair region of the mitochondrial cytochrome c oxidase (COI) marker, using the optimized EPT-specific primers (forward primer fwhF2, reverse primer EPTDr2) from Leese et al. (2021). Next-generation library preparation of three replicate PCR plates was done in the clean laboratory at Eawag (Switzerland, Dübendorf).

We prepared three technical replicate PCR plates per sample (one replicate in each of the 3 plates, at the same position) and followed the protocol of Leese et al. (2021) and Brantschen et al. (2022) with a few adaptions, as described in this section, for all PCR runs (PCR and index PCR). The loading of the extraction and control samples onto a 96-well plate was randomised. Each well was loaded with a total volume of 25 μl: 7.5 μl of DNA sample or control, 0.25 μl of each primer, 12.5 μl of 2x Multiplex MasterMix (Qiagen Multiplex PCR Plus Kit, Qiagen, Hilden, Germany) and 4.5 μl of nuclease-free water (Sigma-Aldrich, St. Gallen, Switzerland). For the PCR negative control (PCRneg), 7.5 μl of nuclease-free water (Sigma-Aldrich, St. Gallen, Switzerland) was added instead of DNA, whereas the extraction negative control (Neg_E) contained sample-free extraction liquid, and the field negative control (Neg_F) contained UV-ed Nanopore water which was filtered through a Sterivex filter (Qiagen, Hilden, Germany) in the field. For the PCR positive controls, three different well-amplifying 142-bp-long sections of non-native invertebrate DNA were used (non-native mosquito (M), non-native moth (P/Pd), non-native Ephemeroptera (S/Sk) and a mix of M and Pd (MPd)).

The initial PCR was run for 33 cycles with annealing temperatures of 64 °C and 58 °C for higher specificity. The PCR program followed a process of initial denaturation for 15 minutes at 95 °C, 10 cycles of 90 seconds of denaturation at 94 °C, primer annealing for 90 seconds at 64 °C, and elongation at 72 °C for 30 seconds. This process was repeated but with a 58 °C annealing temperature for 23 cycles, followed by a final elongation period for 30 seconds at 68 °C. The quality of the PCR product was checked with the QIAxcel Advanced Instrument (Qiagen, Hilden, Germany).

After pooling the PCR product from the 3 technical replicates, the samples were purified using AMPure XP beads (Beckman Coulter, Agencourt, CA, USA) and a magnetic stand. Magnetic beads were added at a 0.7 ratio to improve specificity and selectively remove smaller DNA fragments. The purified PCR products were then washed twice with 200 μl of 80% ethanol on a magnetic stand. To detach the target DNA from the beads, 10 μl of nuclease-free water (Sigma-Aldrich, St. Gallen, Switzerland) were added to the sample. The resulting solution was then pooled in 10-μl aliquots. The purified COI sequences were prepared for indexing by adding 10 μl of a specific combination of Illumina index adapters (Nextera XT Index Kit v2 Set D & A, USA), 5 μl of DMSO (Dimethyl Sulfoxide, PanReac AppliChem ITW Reagents, Darmstadt, Germany), 25 μl of Kappa HiFi HotStart ReadyMix (2x) (Roche, Basel, Switzerland), and 10 μl of sample PCR I on each well.

The index PCR was run for 40 min and 10 cycles, starting with 3 minutes of initial denaturation at 95 °C, and following 10 cycles of 30 seconds at 95 °C, 30 seconds at 55 °C and one minute at 72 °C. Finally, a 10-minute-elongation phase at 68 °C followed. The PCR product quality was assessed using the QIAxcel Advanced Instrument (Qiagen, Hilden, Germany). After annealing the Illumina adapters to the amplicons, a second DNA purification was carried out following the same procedure as previously described, except for two modifications. First, 32.5 μl of nuclease-free water (Sigma-Aldrich, St. Gallen, Switzerland) were added to collect the target DNA from the magnetic beads, and second, the purified supernatant was pooled in 30 μl aliquots. Quality control of the second PCR product was conducted on the TapeStation system 4200 (Agilent, Germany) using the High Sensitivity D1000 ScreenTape (Agilent, Germany), where the reads’ size was captured. DNA quantification was assessed by using the Spark multimode microplate reader (Tecan, Zurich, Switzerland) at the Genetic Diversity Centre (ETH, Zurich, Switzerland). After completion, the samples were pooled and then normalized through a Liquid Handling Station (BRAND, Germany) by targeting a concentration of 10 nanomolars. Finally, the libraries were loaded onto the NovaSeq S Prime Flow Cell and 288 (96 x 3) samples were sequenced at the Functional Genomics Center Zurich (Switzerland) on an Illumina NovaSeq 6000 system (Illumina, San Diego, US) with a 150 bp paired-end read kit v2 with 20% Phi-X spike-in.

### Sequence assignment

Sequences were first demultiplexed into samples using Cutadapt (v.2.10) with the paired adapter mode (Martin 2011). PCR replicates were then demultiplexed, resulting in three distinct replicates per sample. Cutadapt was further utilized for filtering and quality control: non-matching sequences (lacking the primer sequence) were removed and reads shorter than 50 bp were excluded from the analysis. Both the forward and reverse reads (paired reads) were trimmed to a length of 130 bp due to observed quality drops at the end of Illumina NovaSeq reads. Any reads containing ‘N’ were discarded, and those with a maximum expected error greater than 2 were also removed. Furthermore, reads with quality scores equal to or less than 2 were truncated. Sequence error rates were estimated using DADA2 (v.1.16.0), a tool for sequence correction and denoising (Callahan et al. 2016). Following this, a second round of dereplication, sample inference, and merging was performed using DADA2. In this latter step, paired-end reads were combined into single reads. For quality assurance, the DADA2’s BimeraDenovo algorithm was applied to identify and remove chimeras. Taxonomic assignments were achieved through the implementation of the SINTAX algorithm within USEARCH (v.11.0.667, Edgar (2010)). Taxonomy was assigned against the DNA reference database for macroinvertebrates (Leese_Midori_247 COI reference database; Keck and Altermatt (2022)).

Since the expected sequence length is 142 base pairs, only sequences between 137 bp and 147 bp were included in further processing. PCR replicates were pooled, and rare sequences with 10 or fewer occurrences were filtered out. Then, all sequences that had at least 10% of their occurrences in negative or positive control samples were removed, in order to filter out potential contamination. Thereafter, the highest taxonomic resolution for each sequence with a confidence threshold of 0.8 or higher was selected. Lastly, reads corresponding to the same taxon (i.e., species) and PCR replicates were pooled together. Sequencing artefacts (oversaturated reads in two samples) were removed prior to conducting analyses, as they do not accurately represent the real data.

### Statistical analysis

All analyses were performed on presence/absence data at the species level and were run separately for each of the four orders. Presence of a given species was inferred if at least one read for the corresponding sequence was outputted by the bioinformatic pipeline. Site 16 was excluded from all the analyses due to a localized environmental pollution event that occurred some weeks before the start of the sampling campaign and that considerably reduced the size of the community therein, as revealed by our data. We used the software R version 4.3.3 (R Core Team 2024) for statistical analyses and the package *ggplot2* for result visualizations (Wickham 2016).

We used four different approaches to assess complementary facets of phenological signatures in the insect communities of the Necker River. First, we aimed to understand whether the community compositions detected by field replicates were more similar to each other than to those sampled at different time points and/or sites, and to assess whether pairwise differences were more strongly driven by spatial or temporal dynamics. To this end, we performed every possible pairwise comparison of our sampled communities belonging to each of the four investigated orders using the Jaccard index assessed via the *beta.pair* function from the *betapart* package (Baselga et al. 2023). We then categorized the comparisons into four groups according to whether the samples shared the same spatial and/or temporal identity (i.e., they originated from the same site and/or time point). We thus had pairwise comparisons between samples that were taken at a) the same site and time point (“replicate dissimilarity”), b) the same time point but different site (“spatial dissimilarity”), c) the same site but different time point (“temporal dissimilarity”), d) different site and different time point (“spatiotemporal dissimilarity”). Because of the interdependency of Jaccard indices (that is, each sample contributes to Jaccard indices within all of the four groups, depending on the other sample to which it is compared), we refrained from assessing significance of our results via a classical statistical test. Conversely, we designed the following permutation test: for each tested permutation *i* (*i* = 1, …, 10^4^), we shuffled the realized Jaccard index values across all pairs and reassigned them to the four group labels (with each group retaining the same number of pairs as in the original dataset); we then calculated the median Jaccard index *m_i,g_* within each group *g* (*g* = 1, …, 4); we finally compared, for each group *g*, the median Jaccard indices from the original (non-reshuffled) dataset *M_g_* to the so-obtained distribution of *m_i,g_*. Significance at the *x*% level was attributed to the groups whose *M_g_* lied outside of the central (equal-tailed) (100 - *x*)^th^ percentile of the distribution of *m_i,g_*. This is equivalent to assessing whether the median Jaccard index within each group is significantly different than the grand median Jaccard index obtained by pooling all groups.

Second, we wanted to assess whether community changes across time followed specific trajectories. We therefore computed pairwise Jaccard indices across time points and we visualized the findings in 2D space through a Principal Coordinate Analysis (PCoA). To test for statistical differences of the temporal changes, we performed a permutational multivariate ANOVA (PERMANOVA) with 999 permutations using the *adonis2* function of the package *vegan* (Oksanen et al. 2024).

Third, the observed community changes could be driven by either gains or losses in species detectability. To understand which of these two processes underlies the observed community changes across time, we computed temporal beta-diversity indices (TBIs) using the *TBI* function of the *adespatial* package (Dray et al. 2021) and we decomposed the observed changes into gains or losses of species detection. For this approach, we pooled the five field replicate samples to have only one value per site and per time point. A species was considered present at a site and time point if presence was detected in at least two of the five field replicates. For each site, we computed TBIs between the first time point and all successive time points (i.e., the first sampling event was used as a baseline to assess changes). Once again, we used the Jaccard index as dissimilarity method and 999 permutations.

Fourth, we wanted to understand whether our eDNA sampling strategy could replicate temporal signatures of aquatic insect presence that are found through more traditional sampling methods, and hence provide information on when species are most likely to be found. To do this, we extracted observation presence-only data for the four orders from the Global Biodiversity Information Facility (GBIF). The data was constrained to observations recorded between 1980 and 2023 in Switzerland, including only observations listed as “HUMAN_OBSERVATION” (GBIF.org). We also excluded records from the region south of the Alps (i.e., Ticino canton and the Mesolcina valley, due to the strong environmental and biogeographic differences to the study area), taxa for which fewer than 50 observations were recorded, and records taken from November to March. We summarized both GBIF and eDNA data in monthly “frequency of observations” metrics: for GBIF, we took the sum of the observations within each month, whereas for eDNA we took the sum of presences within a time point (thus ranging from 0 to 125), which represented its associated month. For the month of August, for which we had two sampling events, we took the average sum of presences between the two sampling events. We excluded species whose presence in the eDNA data was always below 10 or always above 115 across all months (as this would imply the species was almost always either missing or present, with no temporal trends to track). We then rescaled the eDNA data to represent a frequency of observation (values 0–100%) and subsequently rescaled the GBIF data so that, for each species, the highest monthly value would match the highest monthly frequency of observation found in the eDNA data. Finally, we performed Pearson correlation analyses to compare monthly frequency of observations between the GBIF and the eDNA data.

## RESULTS

We found that the number of species for which we recorded at least one presence within our investigation was similar for Ephemeroptera (n = 35), Plecoptera (n = 45) and Trichoptera (n = 37), but vastly higher for Diptera (n = 1086).

Pairwise analyses of all the samples revealed that field replicates were on average markedly more similar to each other than to other samples (Figure 2), that is, they had lower Jaccard dissimilarity scores (mean number of comparisons per order: 1912 ± 8 SD; the slight difference in number of comparisons across orders is due to a few samples having null communities for a given order). At the other extreme, samples that shared neither space nor time showed the highest dissimilarity scores (n comparisons = 386120 ± 197 SD). The dominance of space or time in driving dissimilarity slightly differed across orders: dissimilarities were more strongly driven by space for Ephemeroptera, Plecoptera and Trichoptera (n = 55130 ± 36 SD), and by time for Diptera (n = 16786 ± 12 SD). Our permutation test revealed that median Jaccard indices within all groups and all orders were highly significant (p < 0.001).

**Figure 2.**
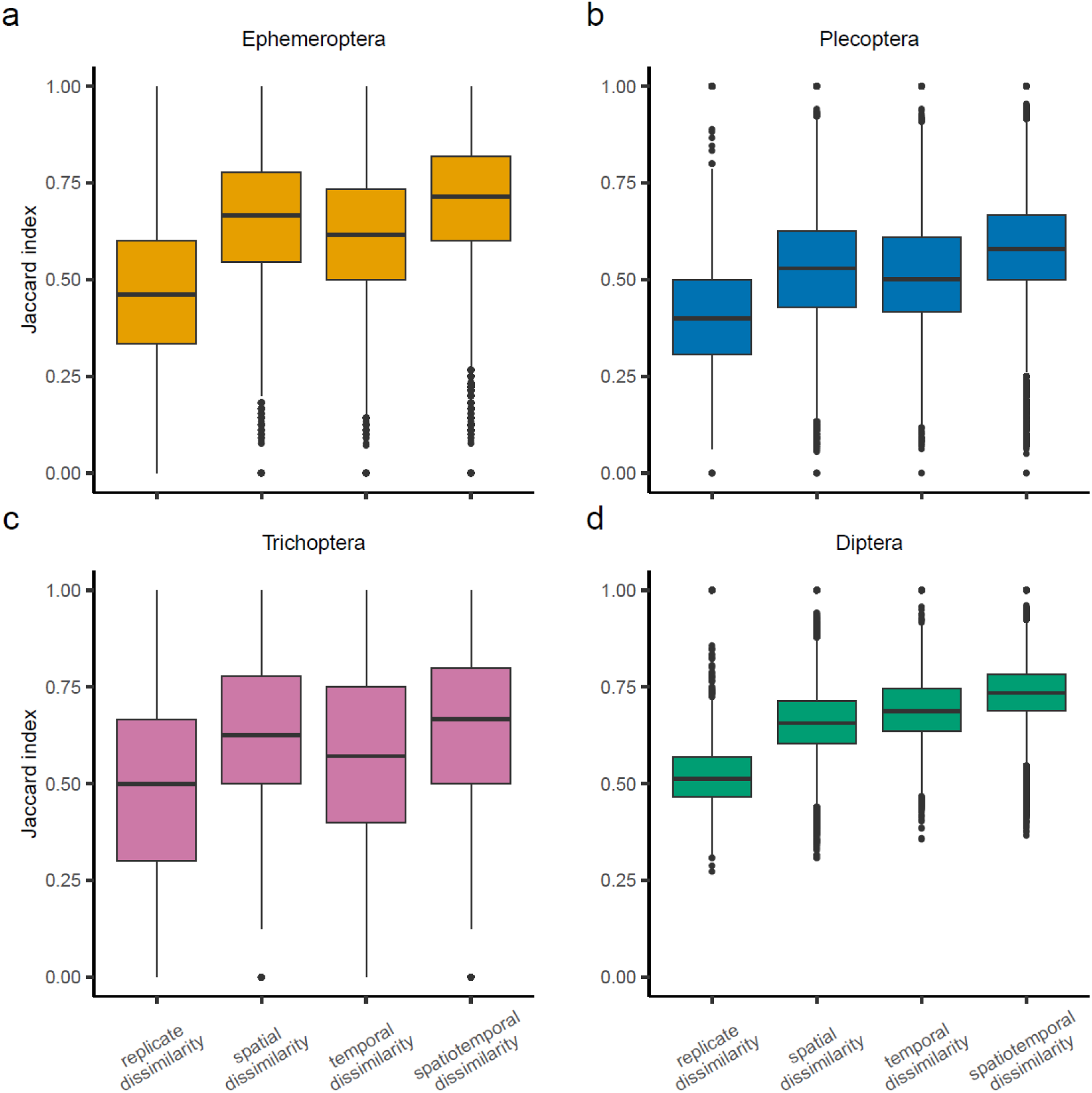
Jaccard indices of dissimilarity obtained by computing pairwise comparisons across all samples and categorized according to the type of dissimilarity investigated (see Methods). Analyses were run separately for the four orders of interest (a–d) and community data was investigated at the species level.

Our PERMANOVA analyses found that the communities of all four orders investigated significantly shifted across time (Ephemeroptera F_7, 926_ = 19.14, p > 0.001; Plecoptera F_7, 931_ = 23.89, p > 0.001; Trichoptera F_7, 913_ = 8.42, p > 0.001; Diptera F_7, 931_ = 29.77, p > 0.001). This shift had a clear boomerang-like trajectory in Ephemeropteran and Dipteran communities (Figure 3a and 3d). The same trend, although less marked, was visible for the order Plecoptera (Figure 3b), whereas the temporal shift detected in Trichopteran communities had a less clear trajectory (Figure 3c).

**Figure 3.**
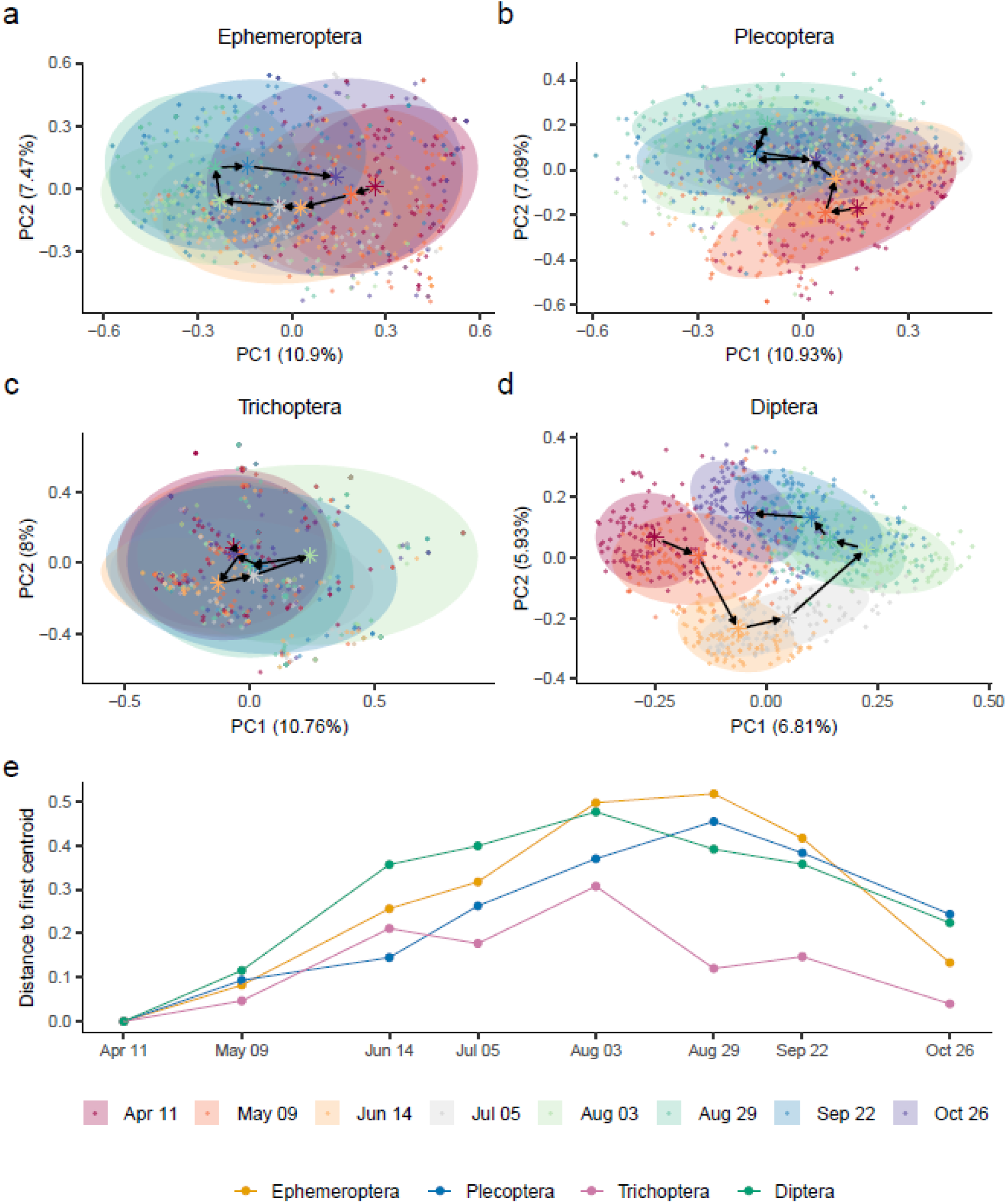
Principal Component Analysis (PCoA) showing the community changes across time for the four major orders investigated (a-d). Points represent the value assigned for each site and replicate at the given week number, ellipses indicate the associated 80% confidence interval, star symbols represent the centroid of the community for that timepoint, and the arrows highlight the movement of the centroid across time. (e) Pairwise distances between the first centroid and the subsequent centroids for all the orders.

The Temporal Beta Index (TBI) analyses revealed similar patterns, but further highlighted differences across orders regarding the nature of the temporal changes. The magnitude of detected community changes—with respect to the first sampling event of April 11—peaked in late August for both Ephemeroptera and Plecoptera, but while changes were driven by gains in species detection for the former, they were primarily due to losses in detection for the latter (Figure 4a and 4b). Changes in Trichoptera were split between gains and losses in species detection throughout spring and early summer, but later in the year were driven by losses (Figure 4c). On the contrary, changes in Diptera were clearly due to gains in species detection early on, while in later summer and fall gains and losses in detection were overall balanced (Figure 4d).

**Figure 4.**
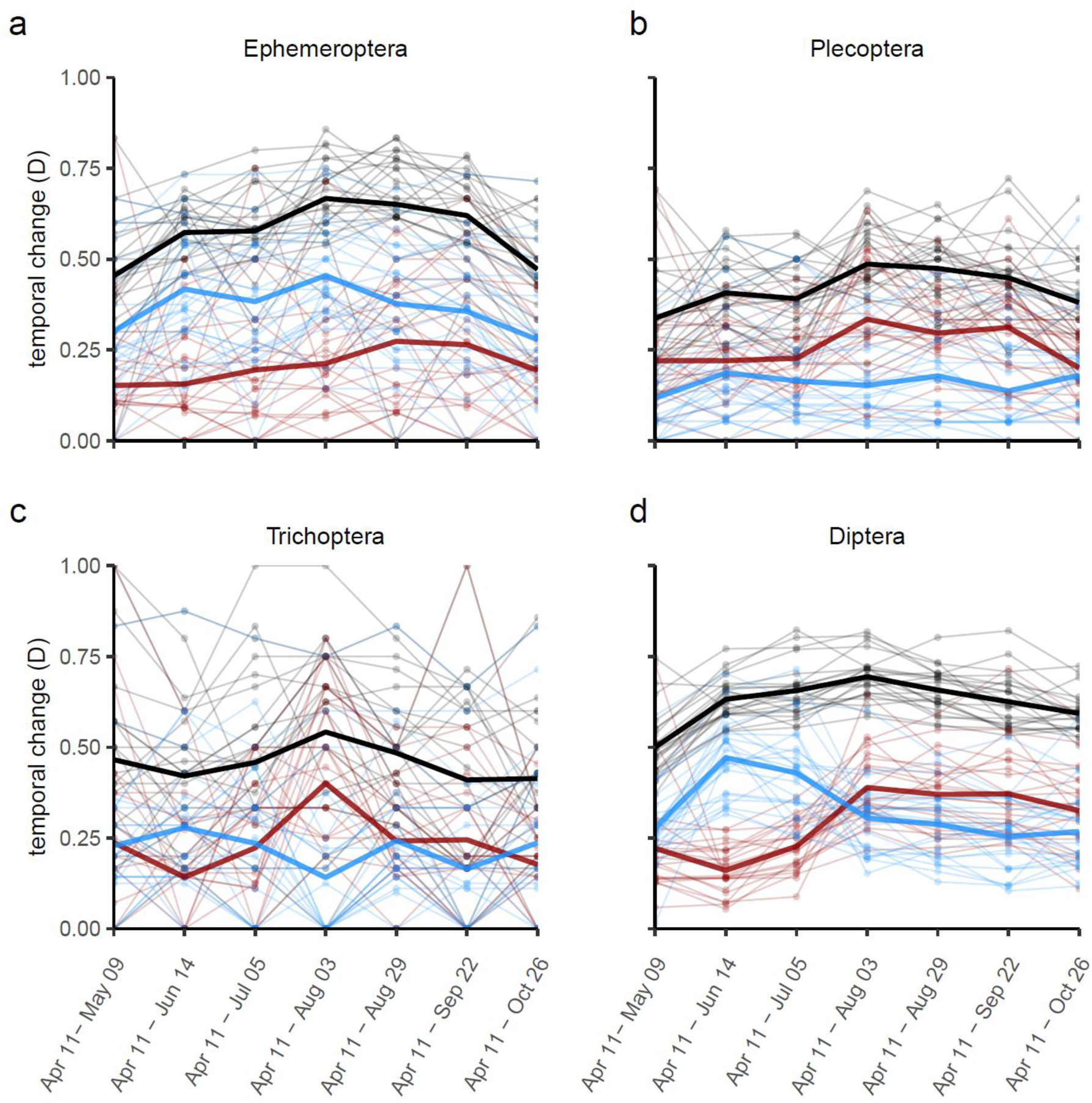
Temporal Beta Index (TBI) for the four major orders investigated (a-d), comparing communities between the first time point sampled (April 11) to all subsequent time points. The total change in community (black lines) is the sum of its components, namely, the gains (blue lines) and the losses (red lines) in species detection. Individual lines and points indicate site-specific values, whereas thicker lines indicate the overall mean at the river level.

We found a total of 64 species that respected our filtering criteria and that could be contrasted between our eDNA data and the GBIF database (Ephemeroptera: n = 23; Plecoptera: n = 27; Trichoptera: n = 9; Diptera: n = 4). Despite the high number of Diptera species found in the eDNA data, records of Diptera species were much scarcer in the GBIF database compared to the other three insect orders, ultimately resulting in the low sample size of Diptera for our comparisons. Using Pearson correlation statistics, we found that temporal trends of frequency of observations between eDNA samples and the GBIF data had a median correlation of 0.380 (Figure 5). No differences were found between orders (all Tukey post-hoc pairwise comparisons: p > 0.9). A Pearson correlation of 0.5 or higher was found in 39% Ephemeroptera species, 37% of Plecoptera, 33% of Trichoptera and 75% of Diptera species (Supplementary material, Figures S1-S4).

**Figure 5.**
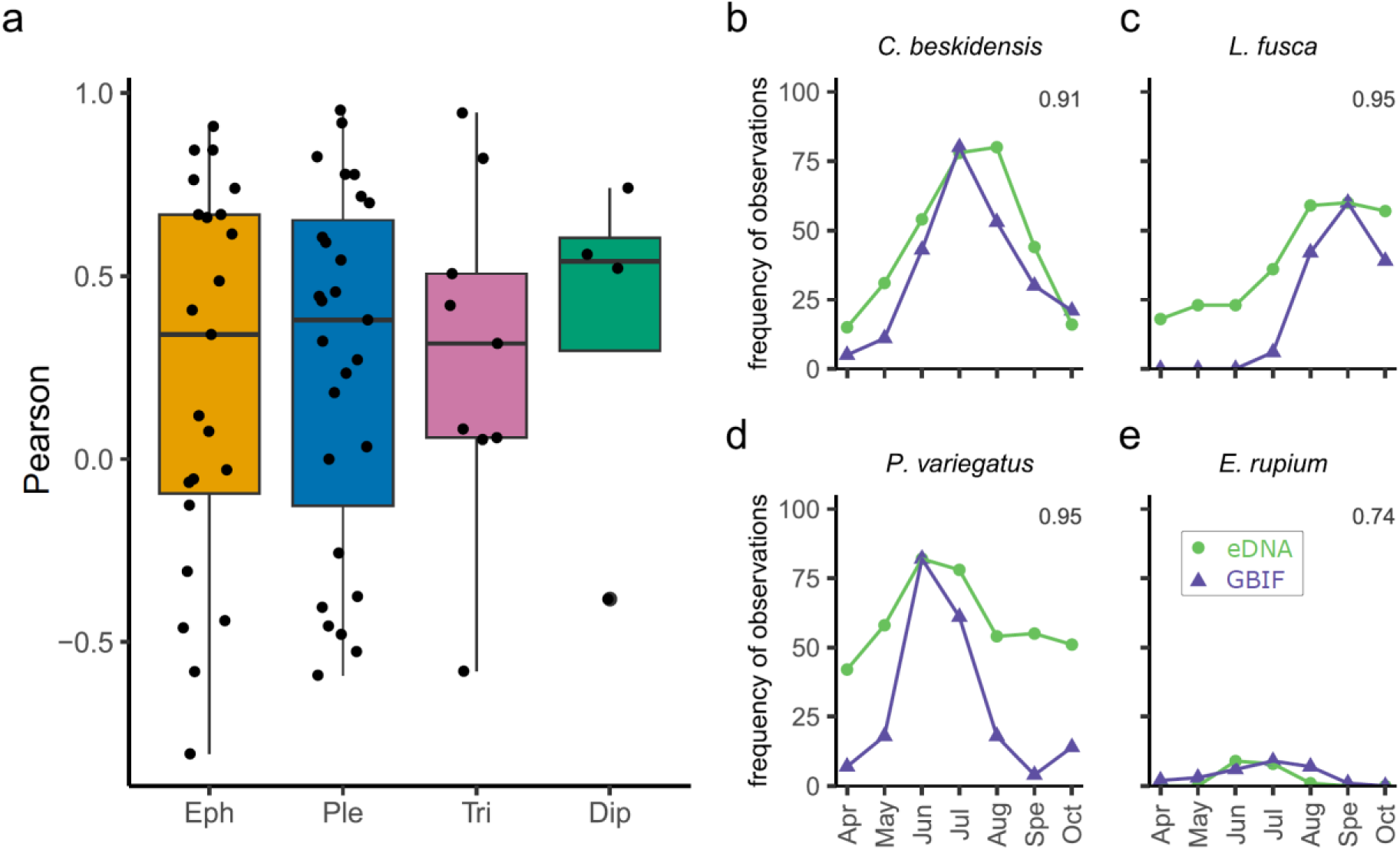
Comparisons of species temporal trends between eDNA samples and GBIF data. Panel a shows the overall Pearson correlation scores between eDNA samples and GBIF data for the four orders. Tukey post-hoc pairwise comparisons revealed no differences between groups. Panels b–e show the frequency of observation obtained with eDNA samples and GBIF data for the four species with the highest Pearson correlation score (indicated in the top right of each panel) per order: (b) *Caenis beskidensis* (Ephemeroptera), (c) *Leuctra fusca* (Plecoptera), d) *Philopotamus variegatus* (Trichoptera) and (e) *Eristalis rupium* (Diptera). The trends for all the investigated species can be found in the Supplementary Material, Figures S1–S4.

## DISCUSSION

We investigated intra-annual dynamics of four major freshwater insect orders (Ephemeroptera, Plecoptera, Trichoptera and Diptera) using roughly monthly samplings of environmental DNA (eDNA) that covered the full extent of a river catchment. We found that both time and space were responsible for structuring communities within all investigated orders. Ephemeroptera, Plecoptera and Diptera communities exhibited cyclic temporal trajectories: from the first sampling in spring, communities diverged over time, reaching the maximum divergence in the mid- to later summer, then returning towards fall to a composition more similar to the one found in spring. These community changes were driven by gains in species detection for Ephemeroptera, losses in detection for Plecoptera and early gains and later losses in detection for Diptera. In the order Trichoptera, we observed a mix of gains and losses in species detection throughout the months, with only hints of temporal cyclic patterns from spring to fall. Overall, we demonstrate that eDNA techniques can be used to reveal intra-annual dynamics of aquatic insects and that for some species these dynamics are reflected in the frequency of observations available on a popular biodiversity database. Given the difficulty of identifying aquatic insects at the species level and the strong phenological signatures of their life history, our findings suggest that eDNA sampling is a viable option to obtain a deeper understanding of the structuring of freshwater communities in time, and, in particular, to capture their pronounced phenology.

As expected, we demonstrated that eDNA is capable of tracking the strong temporal signatures that are typical to aquatic insects. In agreement with the study by Sander et al. (2024), we found for all orders—with the exception of Trichoptera—cyclic trajectories of community changes. These changes are—most likely—not driven by actual shifts in the species composition itself (e.g., as due to by migration, extinction or colonization events), but by changes in the detectability of species due to diverging life history stages and population dynamics. While such cyclic patterns in the detected community structure should therefore be expected in highly seasonal ecosystems with periodic environmental conditions, revealing similar trajectories can be challenging. First, and as mentioned above, the actual species composition of benthic macroinvertebrates might remain constant through the year (Huttunen et al. 2022, Haase et al. 2023), thus changes in the abundance or dominance of species might go undetected. Second, phenological changes in insect communities can be missed if the resolution of the taxonomical identifications is too low or the biodiversity metric being investigated is not refined to detect such dynamics (Šporka et al. 2006). Third, seasonal shifts in detection rates and possibly inferred community structure have been demonstrated using both eDNA methods (Jensen et al. 2021, Blackman et al. 2022) and more conventional methods (Waringer 1996, Cowell et al. 2004, Šporka et al. 2006), but a higher temporal resolution is likely needed to highlight the cyclic transitioning dynamics. The potential of eDNA in capturing temporally highly resolved information was shown for other aquatic environments (Bista et al. 2017, Jensen et al. 2022), and our findings demonstrate that this method is also suited to capturing detailed seasonal signals of the phenology of aquatic insects. Potential caveats of eDNA as a tool to track finely resolved temporal changes could be related to the presence of “old” molecular traces in a water sample, such as due to resuspension of DNA fragments previously adsorbed to the river substrate (Shogren et al. 2017), which might dampen a temporally varying signal. Whether or not this damping effect occurs in our study system, it does not appear so strong as to flatten the temporal dynamics.

We observed the most defined temporal dynamics in Diptera, likely due to the much higher number of species assigned within the order, ultimately allowing the analyses to reveal more clear-cut patterns. Our observation of a switch in mid-summer between the initial gains in species detected and the subsequent loss of species in the signal aligns with past work, which found that the abundance of individuals of Diptera and their detection peaked in early summer just before the emergence of many species (Čmrlec et al. 2013, Sander et al. 2024). Several species of Plecoptera, on the other hand, are known to emerge in earlier spring, therefore species from this order are most commonly detected in (late) winter when they often reach their latest larval instar stages (Waringer 1996, Šporka et al. 2006, Haidekker and Hering 2008). Such phenological trends support our findings of Plecoptera dynamics being underlined by a drop in species detection throughout the sampling campaign in relation to the first sampling event in early April. On the contrary, Ephemeroptera tended to have increased detection of species through spring and summer, and, though this order also displayed cyclic community changes, the overlap between communities at the same time point remained important. In agreement to this result, many species of Ephemeroptera are reported to emerge in summer, but longer emergence duration and the presence of species with multiple generations per year could contribute to diluting this cyclic signal (DeWalt et al. 1994, Waringer 1996, Haidekker and Hering 2008). Lastly, we observed much weaker temporal signals in Trichoptera, which were likely dampened by the lower species richness found. Despite being largely present in this catchment (Mächler et al. 2019), less efficient amplification of Trichoptera sequences and a lower shedding rate of genetic material could hinder the detection of Trichoptera with metabarcoding methods (Leese et al. 2021, Brantschen et al. 2022).

In our work, we utilized field replicates as independent samples to obtain a clearer picture of the uncertainty related to a single eDNA measurement and to allow us to use the presence/absence data as a proxy of frequency of observation. This latter approach appeared promising, revealing temporal dynamics that were, for a selection of species, congruent with those extracted from the Global Biodiversity Information Facility (GBIF). More targeted investigations would be necessary to further validate aquatic insect data on GBIF and assess the possible causes of the discrepancies between our findings and this increasingly popular database. Overall, however, it should be noted that the stochasticity between field replicates was still high. Notably, our Jaccard dissimilarity findings imply that two replicated samples tended to share only about half of their pooled group of species (with median Jaccard indices of about 0.5 across the four orders, see Figure 2). While these dissimilarity values were anyways much lower than the ones obtained for pairs of samples from different time points and/or sites (hence confirming the validity of eDNA as a tool to assess spatiotemporal patterns), they still highlight that assessing communities based on a single sample might result in non-negligible inaccuracies. The importance of filtering large volumes of water in order to obtain samples adequately representing the biodiversity of a water body has been recognized (Altermatt et al. 2023). However, filtering large volumes on a single filter only partially contributes to a complete community assessment, besides being often physically unfeasible due to filter clogging. Indeed, a relevant fraction of the stochasticity in eDNA data is originated in the lab and bioinformatic processing (Burian et al. 2021), and therefore the use of multiple filters related to the same sampling event is a better insurance against possible errors.

Insect communities are core inhabitants of most freshwater rivers across the globe. Although their presence at given sites might be constant through the year, the impact of that presence—that is, their detectability—is expected to vary through the seasons due to the phenological changes in the aquatic insect life history stages and population dynamics. In this study, we demonstrated that eDNA techniques can be used to track the temporal signal of aquatic insect phenologies, helping to shed light on the complex intra-annual dynamics of these biotic communities.

## ACKNOWLEDGMENTS

We thank Samuel Hürlemann for his help with the field data collection and DNA extraction, Heng Zhang and Evelin Pandiamakkal for contributing to the field data collection, and Raphaël Bossart for conducting the library preparation and sequencing. Data were generated in collaboration with the Functional Genomic Center Zurich, Switzerland. EC, AP, NT and LC are supported by the Swiss National Science Foundation Ambizione grant PZ00P2_202010 (to LC) and FA by the Swiss National Science Foundation grant 310030_197410.

## AUTHOR CONTRIBUTIONS

EC, FA and LC conceptualized the study. NT and LC collected the field data. NT conducted the DNA extractions. AP performed the library preparation and sequencing with the support of XA. FK carried out the bioinformatic analyses and taxonomic assignment. EC analyzed the data with the support of XA. EC wrote the first draft of the manuscript. LC supervised the work. All authors contributed to the final version of the manuscript.

