## Appendix for "Tracking the phenology of riverine insect communities using environmental DNA"

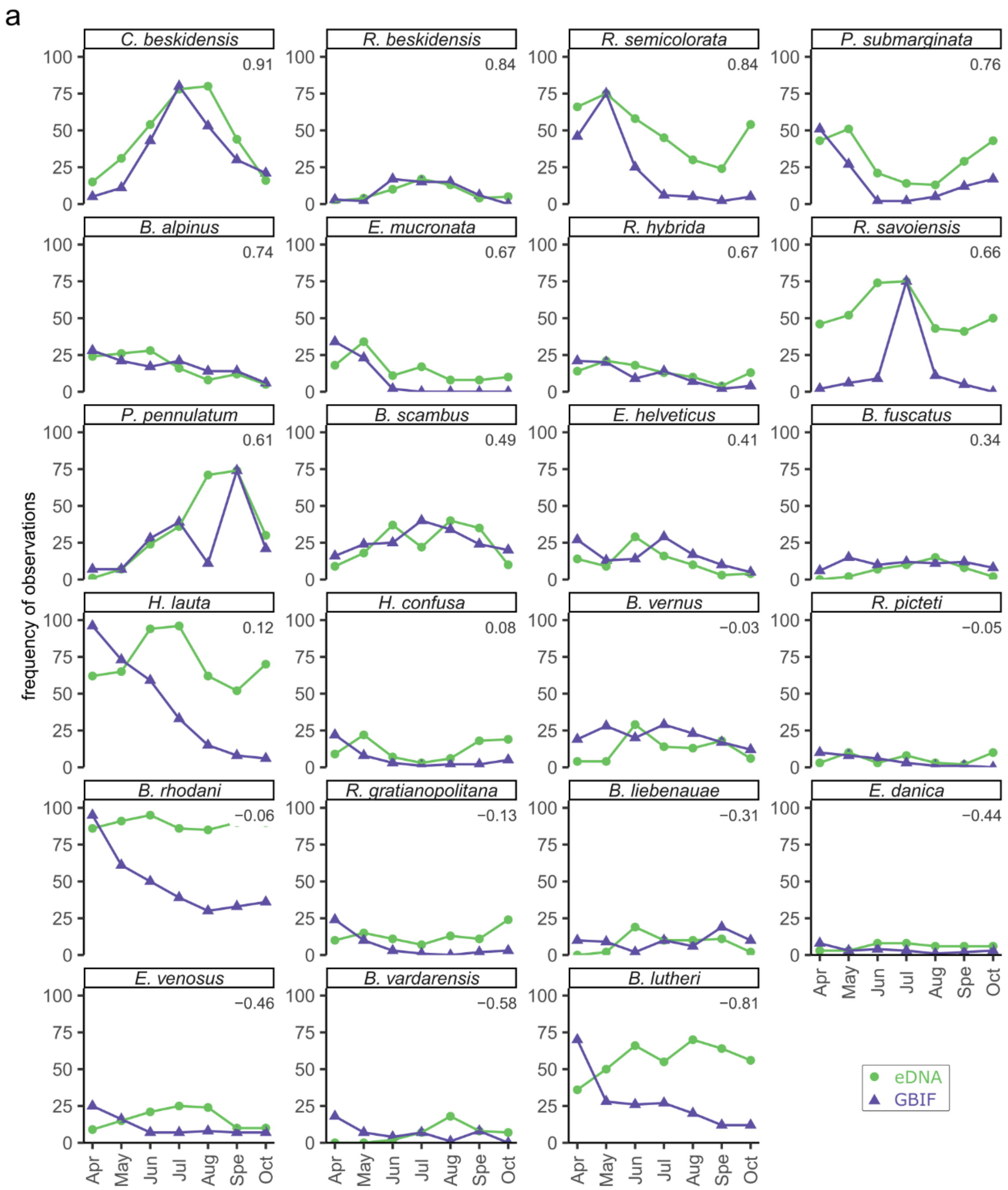


**Figure S1.** Ephemeroptera temporal trends according to eDNA and GBIF data. (a) Comparisons between the frequency of observation found with the eDNA campaign and the one obtained with the GBIF database (and scaled to match the former, see Methods). (b) the associated correlation plots to obtain the Pearson correlation scores (indicated in both panel (a) and (b) at the top of each individual plot). *Continues on the next page.*

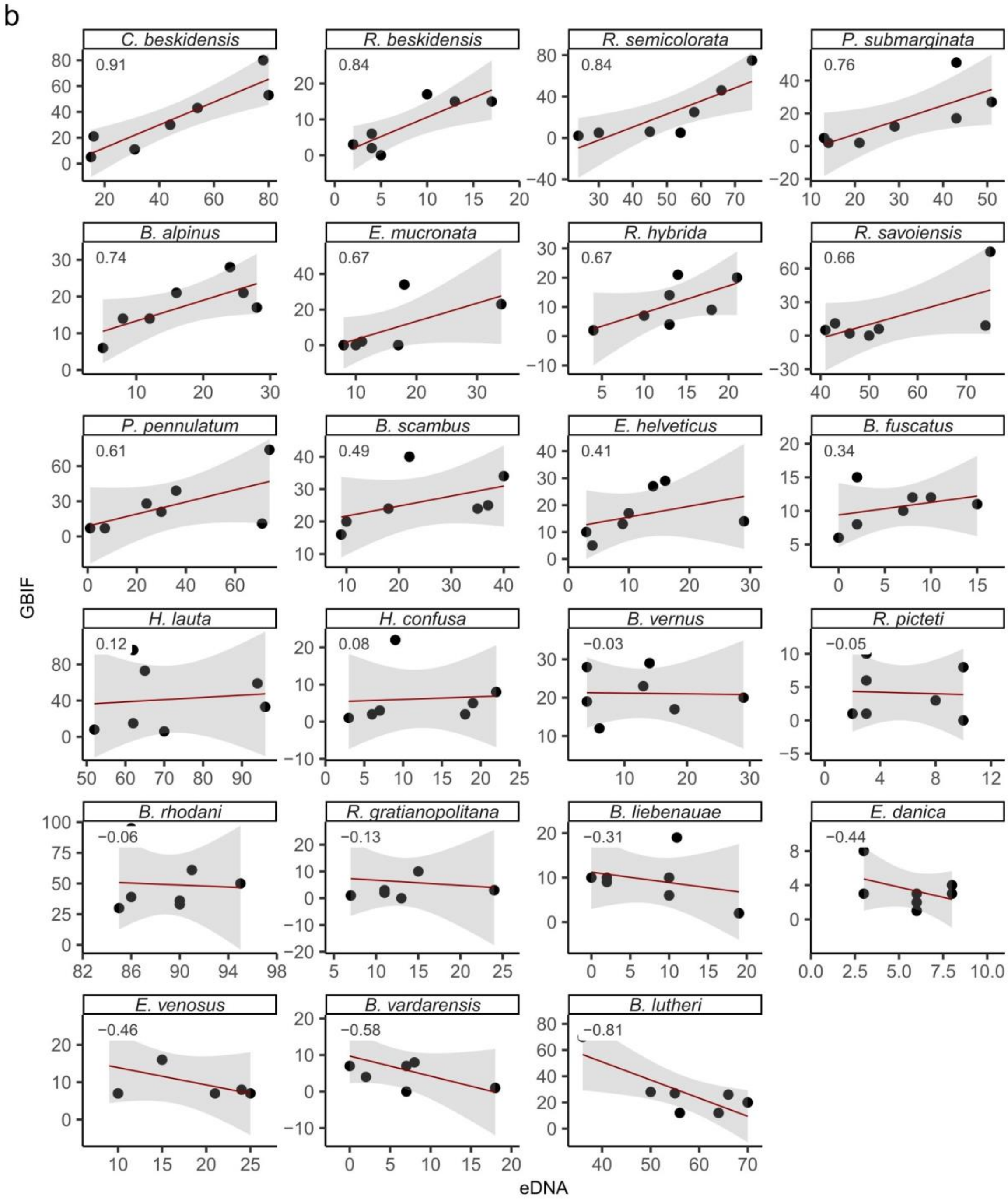


**Figure S1 continuation.** *See caption on previous page.*

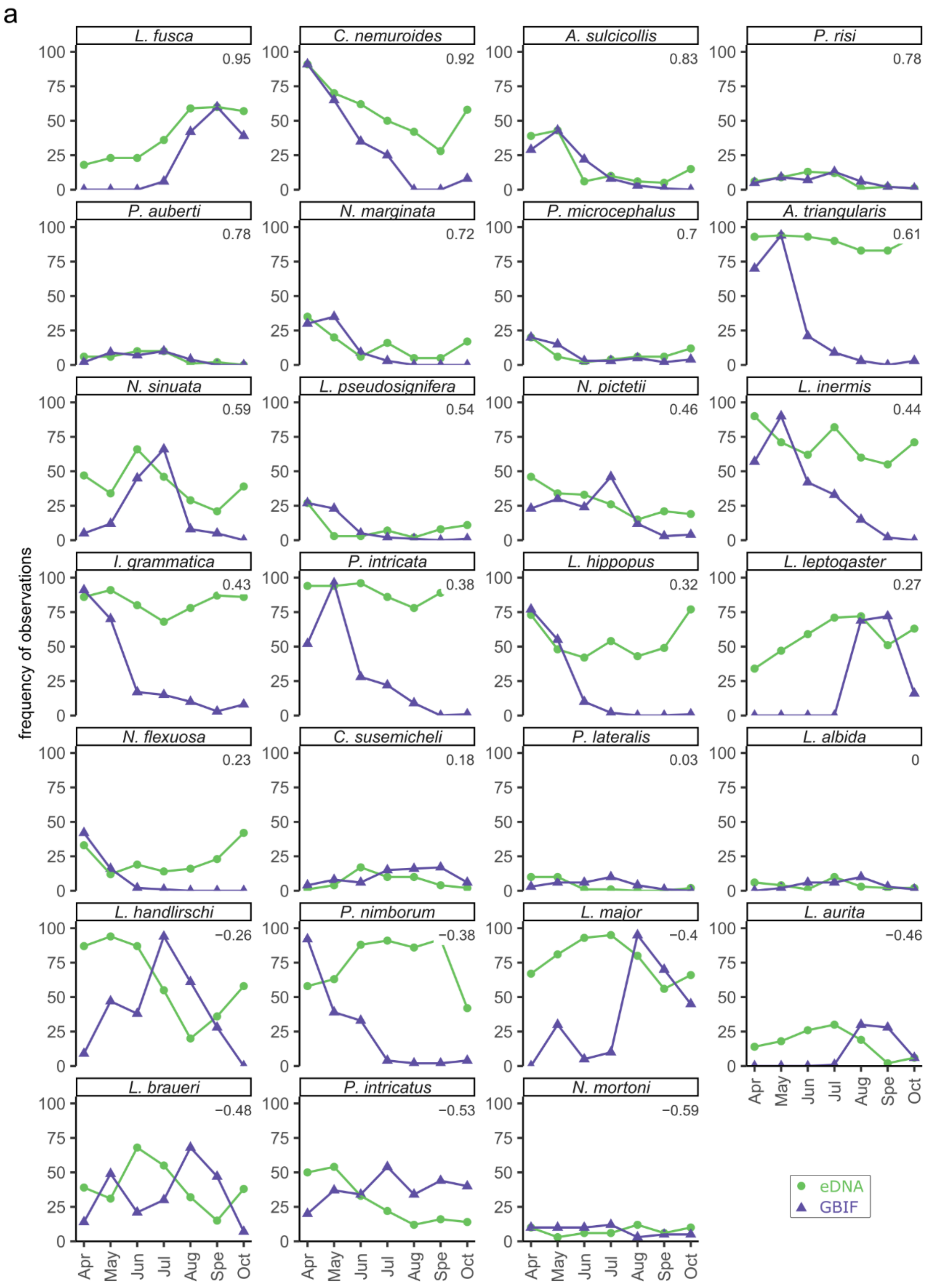


**Figure S2.** Plecoptera temporal trends according to eDNA and GBIF data. See Figure S1 caption for details. *Continues on the next page.*

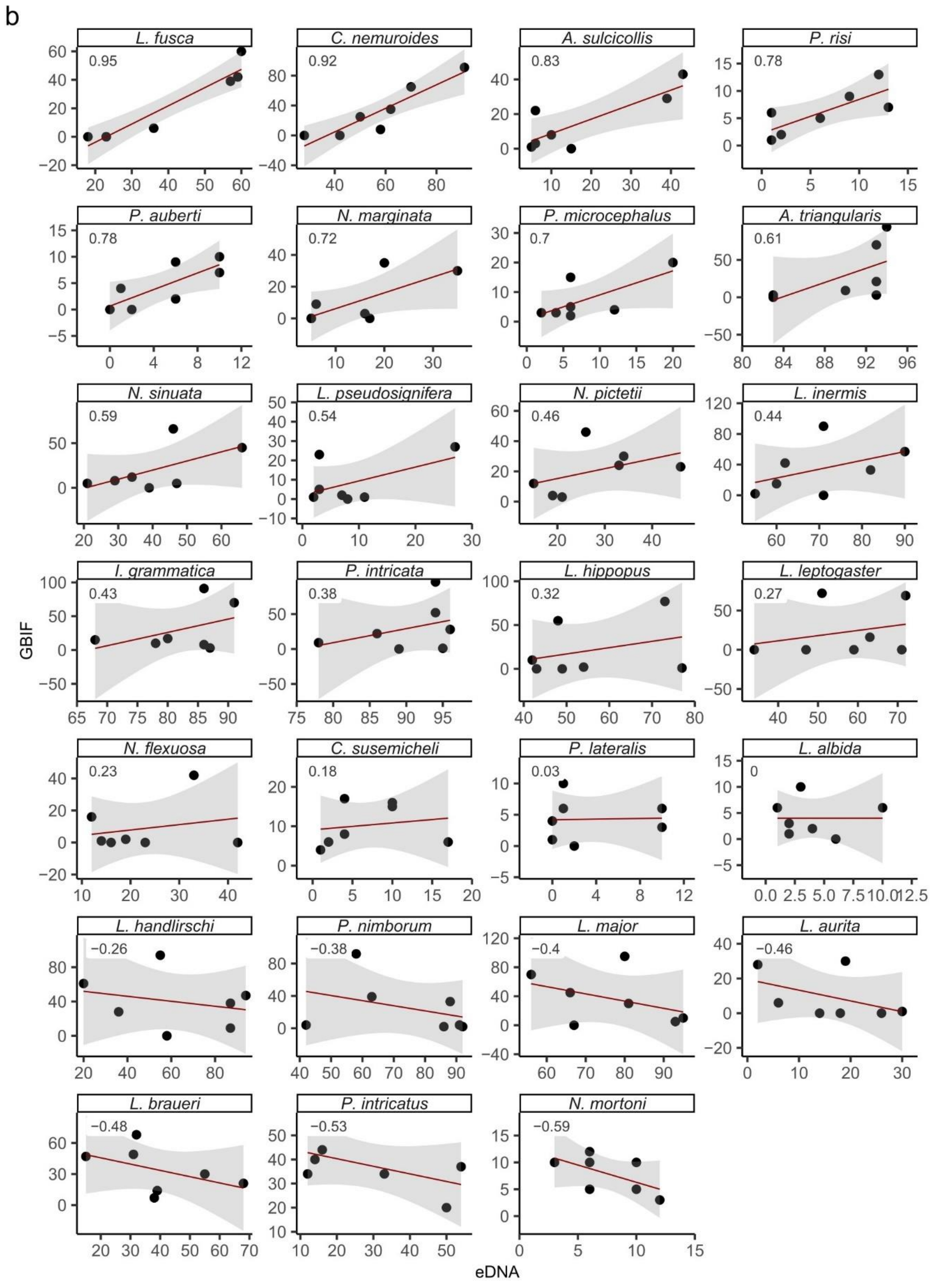


**Figure S2 continuation.** *See caption on previous page.*

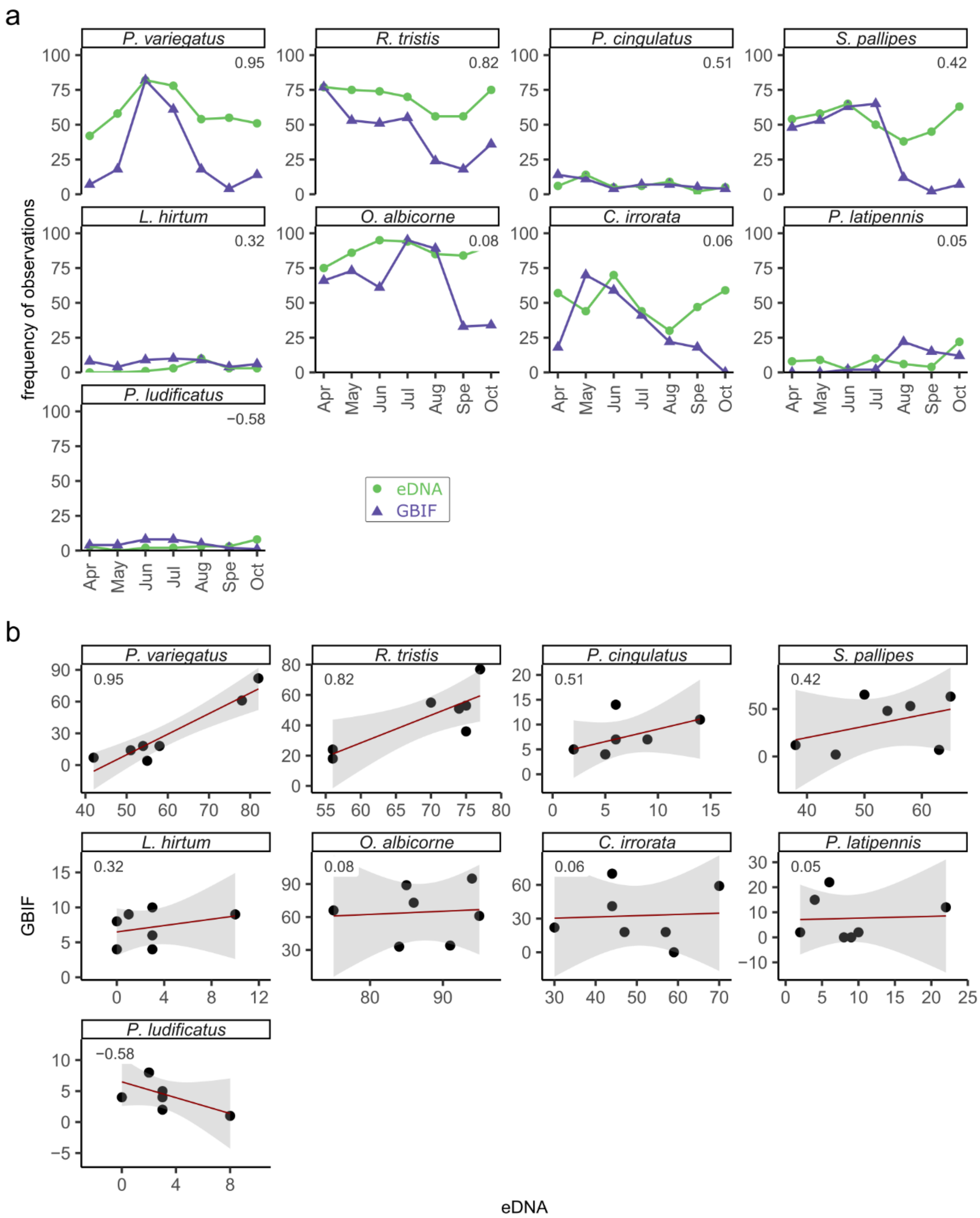


**Figure S3.** Trichoptera temporal trends according to eDNA and GBIF data. See Figure S1 caption for details.

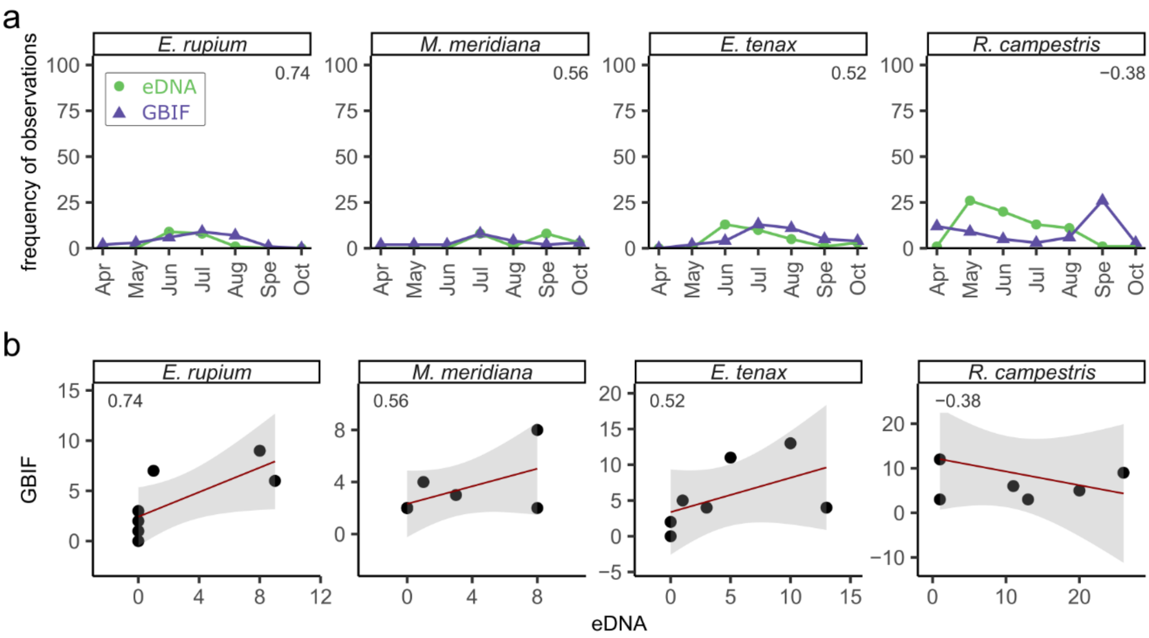


**Figure S4.** Diptera temporal trends according to eDNA and GBIF data. See Figure S1 caption for details.
